# Integrative Analysis of Metabolomic and Transcriptomic Profiles Uncovers Biological Mechanism of Feed Efficiency in Pigs

**DOI:** 10.1101/2020.05.21.108050

**Authors:** Priyanka Banerjee, Victor Adriano Okstoft Carmelo, Haja N Kadarmideen

## Abstract

Feed efficiency (FE) is an economically important trait. Thus, reliable predictors would help to reduce the production cost and provide sustainability to the pig industry. We carried out metabolome-transcriptome integration analysis on 40 purebred Duroc and Landrace male uncastrated pigs to identify potential gene-metabolite interactions and explore the molecular mechanism underlying FE. To this end, we applied untargeted metabolomics and RNA-seq approaches to the same animals. After data quality control, we used a linear model approach to integrate the data and find significant differently correlated gene-metabolite pairs separately for the breeds (Duroc and Landrace) and FE groups (low and high FE) followed by a pathway over-representation analysis. We identified 21 and 12 significant gene-metabolite pairs for each group. The valine-leucine-isoleucine biosynthesis/degradation and arginine-proline metabolism pathways were associated with unique metabolites. The unique genes obtained from significant metabolite-gene pairs were associated with sphingolipid catabolism, multicellular organismal process, cGMP, and purine metabolic processes. While some of the genes and metabolites identified were known for their association with FE, others are novel and provide new avenues for further research. Further validation of genes, metabolites, and gene-metabolite interactions in larger cohorts will help us to elucidate the regulatory mechanisms and pathways underlying FE.

## 1. Introduction

Feed represents about 60%–70% of total pork production costs in modern pig production. Thus, to decrease costs and increase profitability, it is pivotal to identify feed efficient (FE) animals [1]. However, due to the polygenic architecture of FE, individual pigs in a herd exhibit considerable variation in FE despite belonging to similar genetic background and environment [2]. Considering this variation, different methods have been proposed and widely used to measure the FE, including feed conversion ratio (FCR) and residual feed intake (RFI) [1,3]. FCR is the ratio of feed intake (FI) per unit body weight gain and is affected by many factors such as breed, sex, diet, and environmental conditions [4,5]. Pigs with low FCR are considered high FE and vice-versa. RFI estimates the difference between actual and expected FI predicted on production traits as average daily gain (ADG) [6]. Since FCR considers both FI and weight gain, and FCR is also one of the critical predictors of FE, it suggests that feed efficient pigs may possess different physiological-biochemical profiles compared to the inefficient ones [2].

Based on the advances in omics technologies, several approaches have been adopted to shed light on the genetic mechanisms underlying FE in pigs. Among these omics technologies, transcriptomics and metabolomics have helped to elucidate the molecular basis of FE. While transcriptomics allows us to have a transcriptional snapshot of genes underpinning the phenotype under investigation, metabolomics helps to bridge the gap between genotype and phenotype. Recently, an increasing number of metabolomic studies have reported the role of metabolites in economically important traits [7], such as meat quality [8,9], pre-slaughter stress [10], and FE [3]. Likewise, several transcriptomic studies have pointed out candidate genes underpinning FE and other related-traits such as immune response, growth, and metabolism in pigs [11–13]. Despite the new insights into molecular mechanisms reported in these studies, these approaches rely solely on a single regulatory layer without taking into account the multiple interactions among the genomic layers.

To overcome the limitations mentioned above and to gain further insights into biochemical aspects of complex traits, data integration analysis has emerged as a fruitful tool, unveiling potential biomarkers via integration of metabolomics and transcriptomics [14]. By the development of analytical technologies for data integration, the assessment of system-wide changes of transcript levels as surrogate measurements of metabolic rearrangements can be widely assessed. Metabolite-transcript co-responses using combined profiling can yield vital information on the complex biological regulation of the trait. Transcriptome-metabolome integration is a powerful combination as the metabolome is affected by the phenotypic measurements to which the global measures of transcriptome can be anchored [15]. Therefore, herein, we integrated data from metabolome-transcriptome approaches to unveil the unique gene-metabolite pairs. To this end, we adopted a two-step framework, as follows: (1) we first employed the numerical integration of gene-metabolite levels to identify gene-metabolite interaction pairs separately for the breeds (Duroc and Landrace) and FE groups (low and high FE) using IntLIM R-package; (2) Next, a knowledge-based integration approach based on pathway over-representation analysis was used to reveal the underlying pathways in each group (breed-specific and FE-specific). To the best of our knowledge, this is the first study of its kind to ever combine high throughput metabolomics data with RNA sequence based gene expression data in pigs to unravel the missing links between genes and metabolites and to shed light on the molecular basis that characterizes the feed efficiency differences between pigs within and across breeds.

## 2. Results

### 2.1 Descriptive statistics and linear model analysis for genes and metabolites

The data on 749 metabolites and 25,880 genes from 40 samples were generated using untargeted metabolomics and transcriptomic approaches, respectively. We utilized data of 405 annotated metabolites (see methodology) for further analysis. For the transcriptomic data generated on 25,880 genes, we analyzed the data for each of the two groups (breed-specific and FE-specific), as described in the methodology. The genes with a gene count < 1 were removed, resulting in 20,233 genes for both the groups. The gene count data for each group (breed-specific and FE-specific) was normalized, and the linear model was fit into the data as given in the methodology. The genes were also screened for their chromosomal information from the Ensembl *Sus scrofa* database. After normalization, removal of values < 0 and obtaining the gene chromosomal location information, 17,726 (breed-specific), and 17,697 (FE-specific) genes were retained in each group. As a quality control for IntLIM, we filtered out genes with the lowest 5% of the variation, which gave 16,839 genes (breed-specific) and 16,812 genes (FE-specific) that were subjected to IntLIM analysis. A schematic representation of the study design and analysis steps are given in Figure 1. We performed the PCA analysis (Figure S1) on the filtered metabolome-transcriptome data, which included 405 metabolites, and 16,839 genes (breed-specific), and 16,812 genes (FE-specific). The results of PCA analysis for the metabolites-genes based on breed and FE groups are shown in Figure S1.

**Figure 1.**
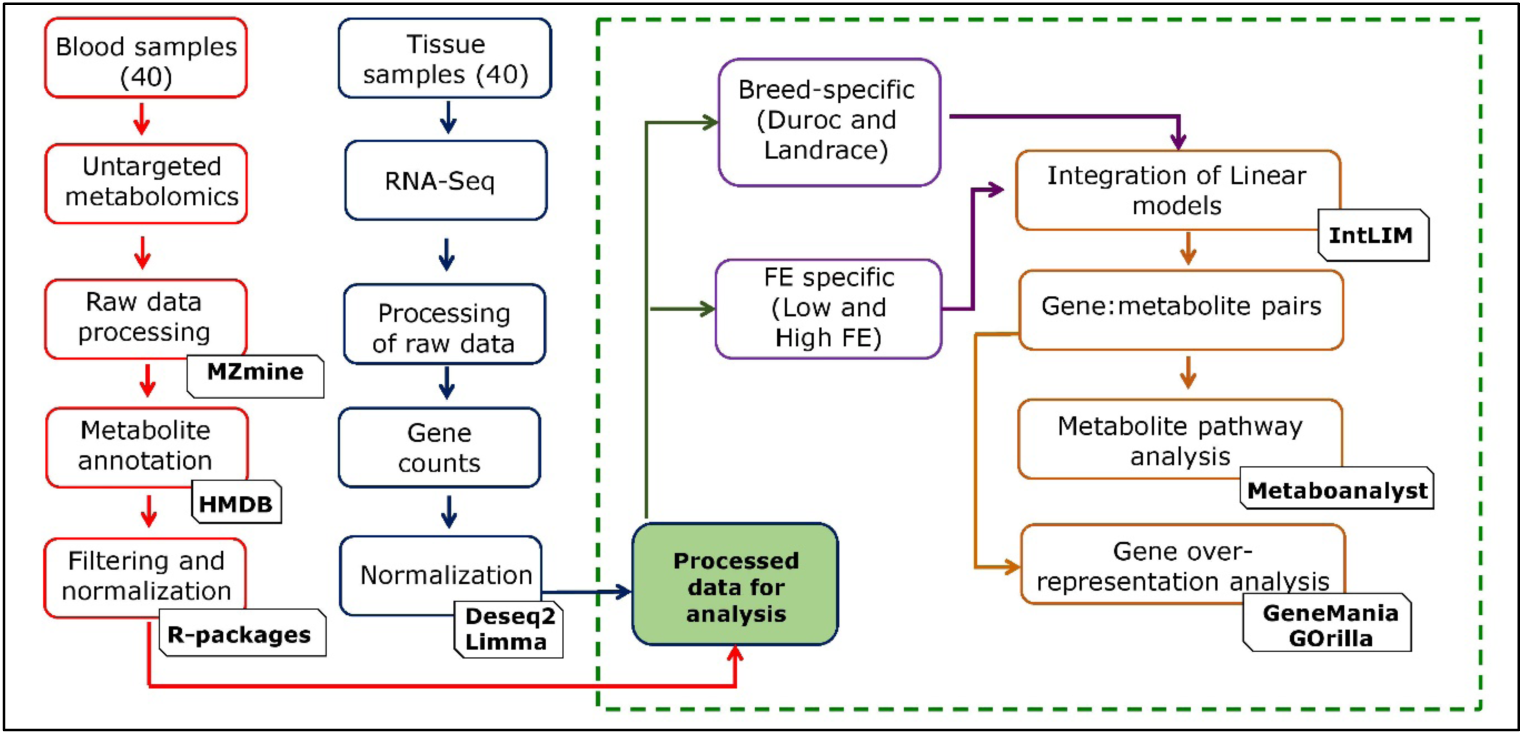
Schematic representation of the study design and analysis steps.

### 2.2 Gene metabolite interaction of breed-specific and FE-specific groups

From the IntLIM analysis, we identified gene-metabolite pairs that have a strong association with respect to breed (Duroc and Landrace) and FE (low and high FE) groups. For the breed-specific groups, all possible combinations of gene-metabolite pairs (6,819,795 model runs) were evaluated, using Duroc and Landrace as the breed-group. Based on this approach, we identified 21 gene-metabolite associations (FDR adjusted p-value < 0.1 and correlation difference effect size > 0.1) (Table 1). Clustering these pairs by the direction of association (positive and negative correlations) within each breed group revealed two major clusters (Figure 2a) in each breed. First, the Duroc correlated/Landrace anti-correlated cluster consists of 7 gene-metabolite pairs (3 unique metabolites and 5 unique genes) with a high positive correlation in Duroc and low or negative correlation in Landrace (Figure 2a). Second, the Duroc anti-correlated/Landrace correlated cluster consists of 14 gene-metabolite pairs (10 unique metabolites and 9 unique genes) with relatively high negative correlations in Duroc and positive correlations in Landrace. The two top-ranked gene-metabolite pairs (ranked in descending order of absolute value of Spearman correlation difference between Duroc and Landrace) in the Duroc correlated and anti-correlated clusters were ENSSSCG00000028124 (*SNRPN*) - Rhodamine B (Figure 2b) and ENSSSCG00000000401 (*GLS2*) - Cystathionine ketimine (Figure 2c) respectively. *SNRPN* and Rhodamine B are positively correlated in Duroc (r = 0.7) but negatively correlated in Landrace (r = −0.5) (Figure 2b). *GLS2* and Cystathionine ketimine are negatively correlated in Duroc (r = −0.9) but positively correlated in Landrace (r = 0.2) (Figure 2c).

**Table 1.** Gene-metabolite interaction pairs from IntLIM for the breed-specific groups.

| Metabolite_name | Ensembl ID | Gene name | Duroc_cor | Landrace_cor | Abs diff.corr | Pval | FDRadjPval |
| --- | --- | --- | --- | --- | --- | --- | --- |
| Rhodamine B | ENSSSCG00000028124 | *SNRPN* | 0.776224 | -0.54242 | -1.31864 | 3.11E-07 | 0.1 |
| L-glutamic acid 5-phosphate | ENSSSCG00000010274 | *SGPL1* | 0.888112 | -0.21456 | -1.10267 | 1.57E-07 | 0.09 |
| Cystathionine ketimine | ENSSSCG00000010274 | *SGPL1* | 0.874126 | -0.18719 | -1.06132 | 2.67E-07 | 0.1 |
| Cystathionine ketimine | ENSSSCG00000040110 | *Novel_gene* | 0.923077 | -0.11385 | -1.03692 | 1.08E-08 | 0.02 |
| L-glutamic acid 5-phosphate | ENSSSCG00000038948 | *ETS2* | 0.895105 | -0.11002 | -1.00512 | 2.66E-08 | 0.02 |
| L-glutamic acid 5-phosphate | ENSSSCG00000040110 | *Novel_gene* | 0.937063 | 0.010947 | -0.92612 | 1.48E-09 | 0.01 |
| L-glutamic acid 5-phosphate | ENSSSCG00000014632 | *FAM160A2* | 0.951049 | 0.229885 | -0.72116 | 1.09E-07 | 0.08 |
| Aloesol | ENSSSCG00000004128 | *ZC2HC1B* | -0.22767 | 0.048714 | 0.276385 | 4.33E-07 | 0.1 |
| Theogallin | ENSSSCG00000026442 | *FAM163B* | -0.29772 | 0.45156 | 0.749284 | 1.69E-07 | 0.09 |
| Fenamiphos | ENSSSCG00000018649 | *Novel_gene* | -0.68652 | 0.093596 | 0.780112 | 3.44E-07 | 0.1 |
| Taraxacolide 1-o-b-d-glucopyranoside | ENSSSCG00000026442 | *FAM163B* | -0.27321 | 0.648057 | 0.921262 | 1.55E-07 | 0.09 |
| Proanthocyanidin a2 | ENSSSCG00000033688 | *ZDHHC22* | -0.63047 | 0.32567 | 0.956144 | 4.20E-07 | 0.1 |
| Fenamiphos | ENSSSCG00000040467 | *Novel_gene* | -0.67251 | 0.288998 | 0.961504 | 2.25E-08 | 0.02 |
| L-glutamic acid 5-phosphate | ENSSSCG00000000401 | *GLS2* | -0.85315 | 0.109469 | 0.962616 | 1.80E-07 | 0.09 |
| Paracetamol sulfate | ENSSSCG00000000401 | *GLS2* | -0.94406 | 0.038314 | 0.98237 | 2.87E-07 | 0.1 |
| L-glutamic acid 5-phosphate | ENSSSCG00000034989 | *LRRTM2* | -0.83916 | 0.15052 | 0.989681 | 2.17E-08 | 0.02 |
| Cystathionine ketimine | ENSSSCG00000034989 | *LRRTM2* | -0.79021 | 0.204707 | 0.994917 | 2.53E-08 | 0.02 |
| Ketoleucine | ENSSSCG00000026442 | *FAM163B* | -0.5289 | 0.466886 | 0.995783 | 1.19E-08 | 0.02 |
| Ganoderenic acid e | ENSSSCG00000034200 | *SEC22C* | -0.85315 | 0.288998 | 1.142145 | 3.47E-07 | 0.1 |
| Cystathionine ketimine | ENSSSCG00000000401 | *GLS2* | -0.93007 | 0.249042 | 1.179112 | 2.92E-09 | 0.01 |
| Ganoderenic acid e | ENSSSCG00000037595 | *Novel_gene* | -0.85315 | 0.449371 | 1.302517 | 2.26E-07 | 0.1 |

Duroc\_cor: Spearman correlation in Duroc; Landrace\_cor: Spearman correlation in Landrace; Abs diff.corr: the absolute difference in correlation between Duroc
and Landrace; Pval: P-value ( $P < 10^{-7}$ was selected as the cut-off value); FDRadjPval: FDR adjusted P-value

**Figure 2.**
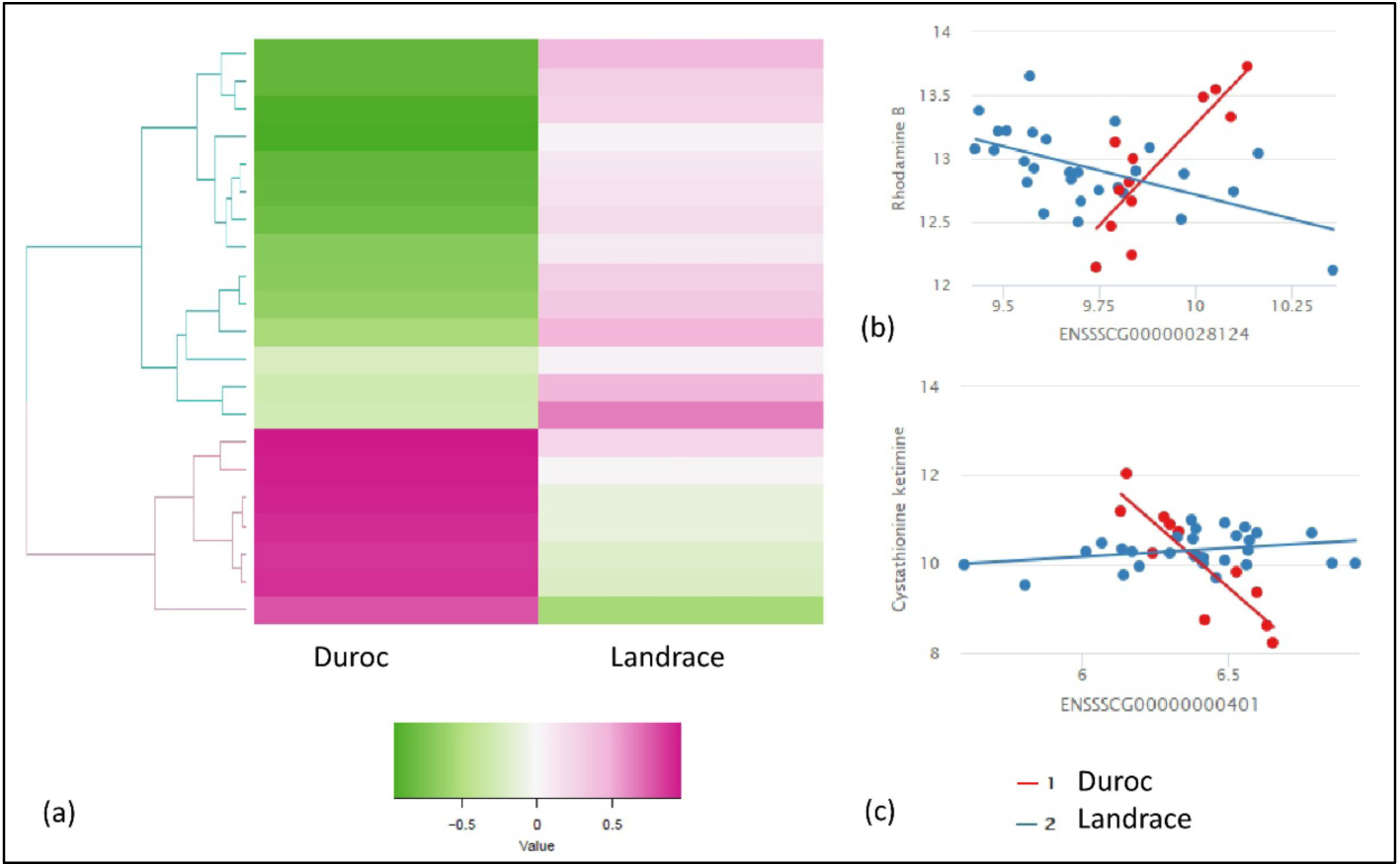
Results of IntLIM applied to breed-specific groups. (a) Clustering of 21 identified gene-metabolite pairs (FDR adjusted p-value of interaction coefficient < 0.1, Spearman correlation difference > 0.1 in Duroc and Landrace breeds, (b) Gene-metabolite difference in ENSSSCG00000028124 (*SNRPN*) - Rhodamine B (FDR adjusted p-value = 0.1, Duroc Spearman correlation = 0.7, Landrace Spearman correlation = −0.5), (c) Gene-metabolite difference in ENSSSCG00000000401 (*GLS2*) - Cystathionine ketimine (FDR adjusted p-value = 0.01, Duroc Spearman correlation = −0.9, Landrace Spearman correlation = 0.2).

Similarly, we used IntLIM for the FE-specific group and evaluated all possible combinations of gene-metabolite pairs (6,808,860 models), with low and high FE as a binary phenotype. With this approach, we identified 12 FE-specific gene-metabolite correlations (FDR adjusted interaction p-value ≤ 0.1, and a Spearman correlation difference > 0.1) involving 8 unique gene and metabolites each (Table 2). The heat-map with gene-metabolite Spearman correlation for low and high FE group showed a clear separation between the two groups (Figure 3a). The high FE-correlated cluster of 12 gene-metabolite pairs (8 unique genes and metabolites with high correlations in high-FE groups) are negatively correlated with the low-FE group. The two gene-metabolite pairs (ranked in descending order of absolute value Spearman correlation difference between high and low FE group) in high-FE correlated clusters were ENSSSCG00000025106 (*THNSL2*) – Pyrocatechol (Figure 3b) and ENSSSCG00000036609 (*TBXT*) – Ketoleucine (Figure 3c), respectively. Both pairs showed a positive correlation in high-FE group (r = 0.6, r = 0.5) while show a negative correlation in low-FE group (r = - 0.7, r = −0.3) (Figure 3b, 3c).

**Table 2.** Gene-metabolite interaction pairs from IntLIM for the FE-specific groups.

| Metabolite_name | Ensembl ID | Gene name | High_cor | Low_cor | Abs diff.corr | Pval | FDRadjPval |
| --- | --- | --- | --- | --- | --- | --- | --- |
| Pyrocatechol | ENSSSCG00000025106 | *THNSL2* | 0.66996 | -0.73284 | -1.4028 | 4.72E-08 | 0.06 |
| 2-pyrocatechuic acid | ENSSSCG00000025106 | *THNSL2* | 0.645257 | -0.68137 | -1.32663 | 7.13E-08 | 0.08 |
| Ketoleucine | ENSSSCG00000017043 | *RNF145* | 0.548419 | -0.75735 | -1.30577 | 2.17E-07 | 0.1 |
| Ketoleucine | ENSSSCG00000025106 | *THNSL2* | 0.498024 | -0.58088 | -1.07891 | 1.19E-07 | 0.10 |
| Theogallin | ENSSSCG00000036609 | *TBXT* | 0.727273 | -0.35049 | -1.07776 | 2.87E-08 | 0.05 |
| Neodiospyrin | ENSSSCG00000029077 | *TUBAL3* | 0.608696 | -0.46814 | -1.07683 | 2.63E-08 | 0.05 |
| Theogallin | ENSSSCG00000025106 | *THNSL2* | 0.4417 | -0.6152 | -1.0569 | 2.52E-07 | 0.1 |
| Proanthocyanidin a2 | ENSSSCG00000008938 | *ENAM* | 0.37954 | -0.60539 | -0.98493 | 1.99E-07 | 0.1 |
| Ketoleucine | ENSSSCG00000036609 | *TBXT* | 0.557312 | -0.3701 | -0.92741 | 8.78E-08 | 0.08 |
| Adrenochrome | ENSSSCG00000038441 | *Novel_gene* | 0.171485 | -0.58088 | -0.75237 | 1.71E-08 | 0.05 |
| Proanthocyanidin a2 | ENSSSCG00000019329 | *U2* | 0.052384 | -0.66176 | -0.71415 | 1.11E-08 | 0.05 |
| Levulinic acid | ENSSSCG00000009250 | *PRKG2* | 0.075117 | -0.54902 | -0.62414 | 1.93E-07 | 0.1 |

High\_cor: Spearman correlation in high FE group; Low\_cor: Spearman correlation in low FE group; Abs diff.corr: the absolute difference in correlation between the high and low FE group; Pval: P-value ( $P < 10^{-7}$ was selected as the cut-off value); FDRadjPval: FDR adjusted P-value

**Figure 3.**
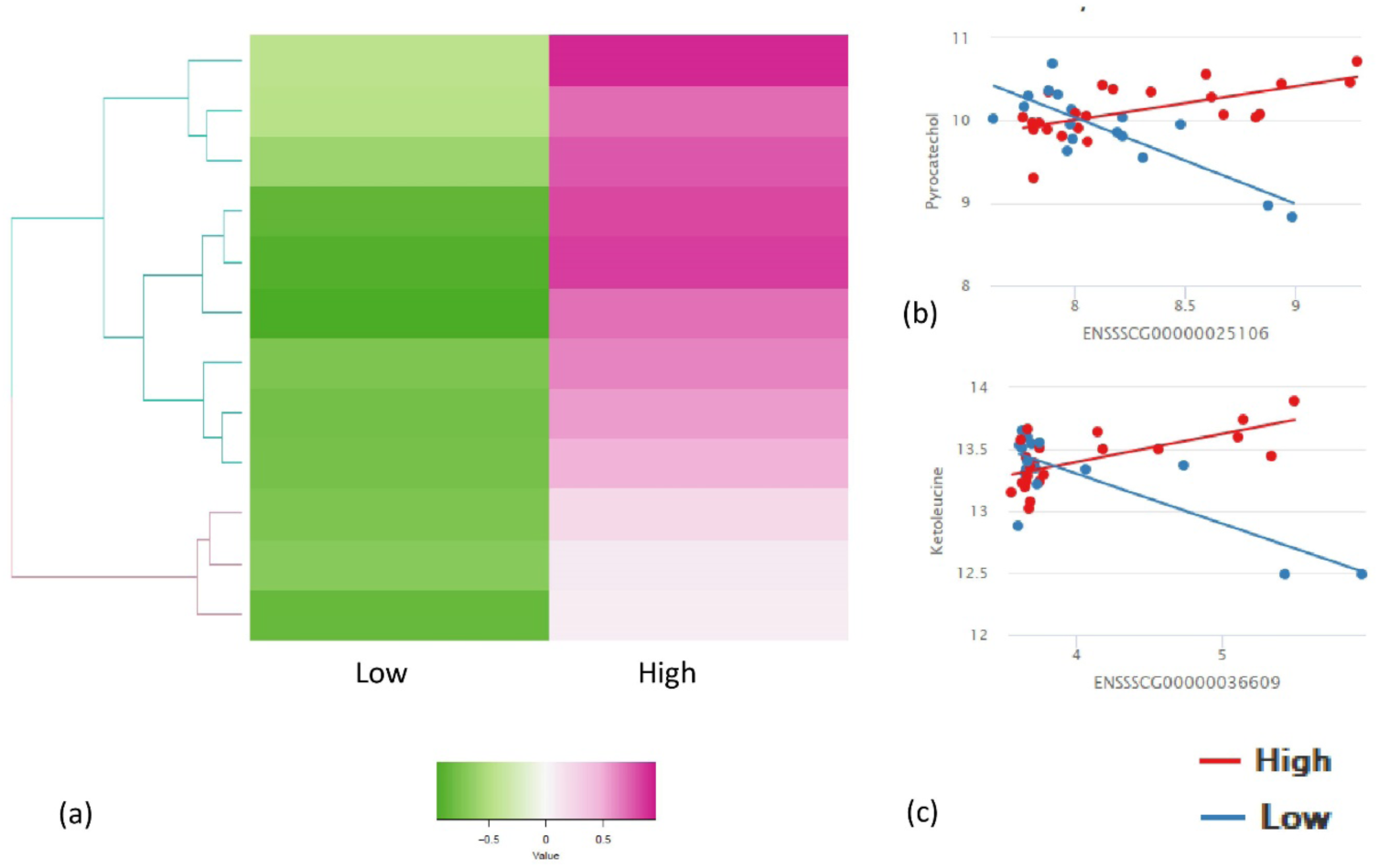
Results of IntLIM applied to FE-specific groups. (a) Clustering of 12 identified gene-metabolite pairs (FDR adjusted p-value of interaction coefficient < 0.1, Spearman correlation difference > 0.1 in high and low FE groups, (b) Gene-metabolite difference in ENSSSCG00000025106 (*THNSL2*) – Pyrocatechol (FDR adjusted p-value = 0.06, High-FE Spearman correlation = 0.6, Low-FE Spearman correlation = −0.7), (c) Gene-metabolite difference in ENSSSCG00000036609 (*TBXT*) – Ketoleucine (FDR adjusted p-value = 0.08, High-FE Spearman correlation = 0.5, Low-FE Spearman correlation = −0.3).

### 2.3 Pathway and gene ontology over-representation analysis

We identified the pathways associated with the unique metabolites in each cluster identified from breed-specific (21) and FE-specific (12) interactions. The three unique metabolites from Duroc correlated/Landrace anti-correlated cluster were associated with arginine and proline metabolism (p-value = 0.02). Furthermore, the ten unique metabolites from Duroc anti-correlated/Landrace correlated cluster were associated with valine-leucine-isoleucine biosynthesis (p-value = 0.01) and valine-leucine-isoleucine degradation (p-value = 0.07) along with arginine and proline metabolism (p-value = 0.07). The eight unique metabolites from high-FE correlated/low-FE anti-correlated cluster were associated with valine-leucine-isoleucine biosynthesis (p-value = 0.01) and degradation (p-value = 0.07). The pathways associated with the metabolites in breed-specific and FE-specific clusters for unique metabolites are given in Supplementary Table S1 - a.

Unique and mappable genes from each group (breed-specific – each cluster, and FE-specific) were screened by using GeneMania to generate a composite functional association network that includes all the evidences of co-functionality. From the breed-specific group, unique genes (4 genes) from Duroc correlated/Landrace anti-correlated cluster (Table 1) mapped to 21 genes based on co-functionality from GeneMania (Table S1 - b). The gene-ontology enrichment analysis of the identified 25 genes (unique genes from Table 1 and co-functional genes from GeneMania) were related to the regulation of hemopoiesis, response to thyroid hormone, and sphingolipid catabolic process (Table S1 – c). Unique and mappable genes (6 genes) identified from Duroc anti-correlated/Landrace correlated cluster (Table 1) were co-functional with 20 genes based on GeneMania (Table S1 – b). The gene ontology enrichment analysis of these 26 genes (unique genes from Table 1 and co-functional genes from GeneMania) revealed the ER to Golgi vesicle-mediated transport and membrane fusion (Table S1 - c) as an enriched pathway. Similarly, from the FE-specific group, unique mapped genes (7 genes) from high-FE correlated/low-FE anti-correlated cluster (Table 2) were co-functional with 20 genes identified from GeneMania (Table S1 – b). The 25 genes (unique genes and co-functional genes) were enriched in the cGMP pathway, purine nucleotide biosynthesis, and phosphorus-oxygen lyase activity pathways (Table S1 – c).

## 3. Discussion

FE is an indicator of efficiency, converting nutrients from the feed to a tissue that is of nutritional and economic significance [16]. Understanding the molecular mechanisms underlying FE will help in the efficient selection of pigs and benefit the pig producers. In the Danish pig industry, Duroc is the most popularly used terminal sires in combination with crossbred Landrace X Yorkshire breeds [17], so the selection pressures for FE in Duroc is higher as compared to Landrace. Thus, in our study, we attempted to identify the gene-metabolite interactions specific to each breed. FE is a complex trait influenced by environmental and health factors and involves many organs. Skeletal muscle, being the largest organ in the body, is an essential location for the metabolism of carbohydrates and lipids [18,19]. It plays a significant role in the utilization and storage of energy acquired from the feed. Many metabolic processes underlie the transport of molecules through the blood, which is the sole way of absorption of nutrients in the body, and blood metabolites are a prime candidate for the study of FE. Thus, understanding the biological processes with the markers obtained from muscle and blood from diverging FE groups will help to optimize the strategies to improve FE and decrease production costs.

A plethora of metabolome and transcriptome studies for FE in pigs are reported [3,9,10,12]. However, to the best of our knowledge, markers from the integration of metabolome and transcriptome in Duroc and Landrace pigs has not been done before. Herein, we unveiled the gene-metabolite relationships that are phenotype dependent. This approach highlighted molecular functions and pathways that are strongly evidenced by the integration study. Evaluating phenotype-specific relationships between metabolites and genes helped to elucidate novel co-regulation patterns that would not be identified by single approaches. In our study, we integrated untargeted metabolomic and transcriptomic data. We used a numerical data integration approach that employed the integration of a linear model (IntLIM package) to predict metabolite levels from gene-expression in a phenotype dependent manner [20].

We attempted to identify the breed-specific and FE-specific gene-metabolite pairs in our study. The PCA analysis showed a difference in the expression of genes in Duroc and Landrace. However, PCA for the FE group did not exhibit significant clusters between groups, which may be due to the small sample size evaluated here. With our current metabolome-transcriptome analysis, we identified 21 gene-metabolite breed-specific pairs and 12 gene-metabolite FE-specific pairs.

In the breed-specific analysis, two clusters were identified between Duroc and Landrace breeds. The Duroc correlated/Landrace anti-correlated cluster identified L-glutamic acid 5-phosphate metabolite with *FAM160A2*, ENSSSCG00000040110, *ETS2*, and *SGPL1* genes; Cystathionine ketimine metabolite with ENSSSCG00000040110 and *SGPL1* genes; Rhodamine B with *SNRPN* genes. A previous study showed the *SNRPN* gene (small nuclear ribonucleoprotein polypeptide N) expression in liver, lung, small intestine, skeletal muscle, heart, kidney, spleen, inguinal lymph nodes and fat in pigs [21]. This study also reported the maternal imprint of the *SNRPN* gene in the skeletal muscle of neonate pigs [21]. Small nuclear ribonucleoproteins and heterogeneous small nuclear riboproteins play roles in nucleolar ribosomal RNA (rRNA) and messenger RNA (mRNA) synthesis in conjunction with spliceosome activity responsible for cleaving on introns from pre mRNA molecule [22]. Furthermore, in a study reported for feed efficiency in broiler chickens, high feed efficiency phenotype exhibited enrichment of ribosome assembly including small nuclear ribonucleoprotein, as well as nuclear transport and protein translation processes than low FE phenotype [23]. In a previous study, *ETS2* gene was reported to be differentially expressed and significantly enriched in cell-survival function in high FE pigs [24]. *ETS2* - L-glutamic acid 5-phosphate gene metabolite pair is identified in Landrace anti-correlated cluster in our study, L-glutamic acid 5-phosphate was also a key metabolite in Landrace as evident from the previous study [3]. Arginine and proline metabolism pathways were associated with the unique metabolites from this cluster (p-value ≤ 0.05). The gene ontology enrichment analysis of the unique genes identified from this cluster with the co-functional genes enriched for multicellular organismal process, sphingolipid catabolic process, regulation of hemostasis, and coagulation pathways (p-value ≤ 0.05).

The Duroc anti-correlated/Landrace correlated cluster identified aloesol - *ZC2HC1B*, Proanthocyanidin a2 - *ZDHHC22*; Fenamiphos metabolite with ENSSSCG00000040467, ENSSSCG00000018649 gene; Ganoderenic acid e metabolite with ENSSSCG00000037595 and *SEC22C* genes; *FAM163B* gene with Taraxacolide 1-o-b-d-glucopyranoside, theogallin, and Ketoleucine metabolites; *LRRTM2* gene with cystathionine ketimine and L-glutamic acid 5-phosphate metabolites and *GLS2* gene with L-glutamic acid 5-phosphate, Cystathionine ketimine, and paracetamol sulfate metabolites. A previous study of transcriptome analysis on feed efficient beef cattle revealed *GLS2* gene as one of the differentially expressed genes reducing hepatic lipid synthesis and accumulation [25]. Valine-leucine-isoleucine biosynthesis and degradation in addition to arginine and proline metabolism pathways were associated with the unique metabolites from this cluster (p-value ≤ 0.05). The unique genes of Duroc anti-correlated/Landrace correlated cluster with the co-functional genes identified enrichment for ER to Golgi vesicle-mediated transport and membrane fusion pathways after gene ontology enrichment analysis (p-value ≤ 0.05).

The unique metabolites identified in the breed-specific clusters were also previously reported in another study for the metabolomic analysis of FE related traits in Duroc and Landrace [3]. The breed-specific unique metabolites such as Aloesol and Ketoleucine affected FE in Duroc [3]. In contrast, Rhodamine B, Taraxacolide 1-o-b-d-glucopyranoside, and ganoderenic acid were underlying testing daily gain (TDG) in Duroc [3]. Theogallin and ketoleucine were involved with TDG and daily gain (DG) in Duroc and Landrace and RFI in Duroc [3]. L-glutamic acid 5-phosphate, cystathionine ketimine, and paracetamol sulfate were associated with FE and RFI in Landrace [3].

In the FE-specific analysis, we found 12 significant gene-metabolite pairs. The gene-metabolite pairs in high-FE correlated/low-FE anti-correlated cluster were *TBXT* gene with theogallin and ketoleucine metabolites; *THNSL2* gene with pyrocatechol, 2-pyrocatechuic acid, ketoleucine, and theogallin metabolites; *TUBAL3* gene with neodispyrin metabolite, *RNF145* gene with ketoleucine metabolite, *ENAM* gene and U2 snRNA with proanthocyanidin a2 metabolite, ENSSSCG00000038441 gene with adrenochrome metabolite, and *PRKG2* gene with levulinic acid metabolite. The pathway analysis with the unique metabolites identified from high-FE correlated/low-FE anti-correlated cluster was over-represented for valine-leucine-isoleucine biosynthesis and degradation pathway. The unique mapped genes and co-functional genes were enriched for the following biological processes: lyase activity, cGMP metabolic process, phosphorus-oxygen lyase activity, and cyclic purine nucleotide metabolic process. Ketoleucine is an abnormal metabolite that arises from the incomplete breakdown of branched-chain amino acids (https://hmdb.ca/metabolites/HMDB0000695). Branched-chain amino acids include leucine, isoleucine, and valine that play a critical role in energy homeostasis in addition to lipid and protein metabolism [26]. A significant role of lipid and protein metabolism from the genes differentially expressed in high-FE pigs have been reported earlier [27]. Ketoleucine is also regulated by branched-chain α-keto acid dehydrogenase. Studies reported that the branched-chain α-keto acid dehydrogenase catalyzes the irreversible oxidative decarboxylation of all three branched-chain keto acids (BCKA) derived from BCAA, *i.e*., α-ketoisocaproate (ketoleucine) [28]. The demonstrated changes in BCKAD activity showed a significant alteration in branched-chain amino acid (BCAA) and protein metabolism during starvation in rats [28]. Gene-metabolite interaction of Ketoleucine with *TBXT, THNSL2*, and *RNF145* gene was identified in our study. *THNSL2* was reported among the top 40 significantly differentially expressed genes of characterized proteins between high- and low-ADG steers from a liver transcriptome profiling of beef cattle [29]. This gene encodes a threonine synthase-like protein. A similar enzyme in mouse is reported to catalyze the degradation of O-phospho-homoserine to a-ketobutyrate, phosphate, and ammonia. This protein also has phospho-lyase activity on both gamma and beta phosphorylated substrates (https://www.ncbi.nlm.nih.gov/gene/55258). This gene has also been associated with abdominal and visceral fat in humans based on GWAS [30]. *RNF145* participates in key signaling pathways of cholesterol homeostasis [31]. *RNF145* expression is induced in response to LXR activation and high-cholesterol diet feeding [31]. Transduction of *RNF145* into mouse liver inhibits the expression of genes involved in cholesterol biosynthesis and reduces plasma cholesterol levels. On the other hand, its inactivation increases cholesterol levels both in the liver and plasma [31]. The unique genes from the high-FE cluster were enriched for the cGMP pathways, similar to the studies reported earlier with feed efficiency related traits with pigs [32] and beef cattle [33]. Cyclic nucleotides 3’,5’-cyclic adenosine monophosphate (cAMP) and 3’,5’-cyclic guanosine monophosphate (cGMP) are ubiquitous intracellular second messengers that regulate multiple physiological functions [34]. *PRKG2* gene, one of the main predictors for cGMP pathways, encodes serine/threonine-protein, which binds to inhibits the activation of several receptor tyrosine kinases and is a regulator of the intestinal secretion, bone growth, and renin secretion (https://www.ncbi.nlm.nih.gov/gene/5593). cGMP pathway was also found to affect feed efficiency related traits in Landrace from a previous study [32]. An interaction of proanthocyanidin a2 with ENSSSCG00000019329 (U2 snRNA) was also identified with low but positive correlation with high-FE cluster while a negative correlation with the low-FE cluster. U2 spliceosomal snRNAs are the molecules found in the major spliceosomal machinery of all eukaryotic organisms and affect their gene expression [35]. U2 snRNA plays a central role in the splicing of mRNA precursors by regulating a dynamic set of RNA-RNA base-pairing interactions [36]. From the previously reported studies, the role of precursor mRNA in gene expression has been established as it removes the intronic sequence from immature RNA, leading to a production of mature mRNA that might differ in function [37]. Regulation of pre-mRNA splicing by nutrients modulates the carbohydrate and lipid metabolism [37]. Altered nutrient utilization in FE-divergent animals revealed that high-FE pigs favor metabolic shifts comprising lipid and carbohydrate utilization when compared to low-FE pigs [27]. U2 interacts with proanthocyanidin a2 in our study. Proanthocyanidin a2 is an antioxidant and has a broad spectrum of biologic properties against oxidative stress [38]. Proanthocyanidin significantly increased activity of antioxidant enzyme such as superoxide dismutase, glutathione peroxidase, and catalase [38]. Role of antioxidant activity with FE was reported earlier in beef cattle as low feed efficient steers had greater superoxide dismutase and glutathione peroxidase activity than the high feed efficient ones, potentially using a greater proportion of energy [39]. Thus, as evident from all these facts, potential role of proanthocyanidin a2 – U2 interaction in high-FE pigs identified in our study might be interesting to explore.

The gene-metabolite pairs identified in our study over-represented some pathways that have been reported to have a role in feed efficiency related traits. Some of the genes identified are novel and were not included in the pathway analysis. Since these gene-metabolite pairs selected have a highly significant correlation with respect to each study group, a detailed study of these genes and metabolites are needed to understand their role in feed efficiency related-traits. Further studies on the identified gene-metabolite pairs may help for the discovery of biomarkers as these significant pairs identified directly reflect the phenotype as revealed by the candidate gene-expression with the downstream metabolite differences in pigs with low and high FE groups.

## 4. Materials and Methods

### 4.1 Data resource and phenotype generation

The pigs used in this experiment were raised at the pig testing station “Bøgildgård” operated by SEGES within Landbrug and Fødevarer (L&F: Danish Agriculture and Food Council). Pigs were *ad libitum* fed and free water supplied. The authors of this study were not responsible for animal husbandry, diet, and care as the testing station is a facility within the Danish breeding program run by SEGES. The initial body weight of the pigs before the testing period was approximately 7kg, followed by a 5-week acclimatization phase. For the phenotypic traits, we calculated FCR and RFI, as reported in our previous study [32]. We considered the same classification of animals in our study as efficient and inefficient (low and high FCR, respectively), as previously reported [32]. The classification was done by selecting pigs that were one standard deviation above or below the mean FCR for each breed as previously reported.

### 4.2 Gene expression profile, metabolite profile, and data analyses

For transcriptome analysis, we collected *psoas major* muscle from 40 purebred male uncastrated pigs from two breeds comprising of 12 Danbred Duroc and 28 Danbred Landrace. The tissue samples were preserved in RNAlater (Ambion, Austin, Texas) immediately post-slaughter and stored at −25 °C until subsequent analysis. Total RNA isolation and sequencing were carried out by BGI Genomics. Paired-end sequencing (100 bp) was performed on the BGISEQ-500 platform after Oligo dT library preparation. Read quality control, mapping, and gene counts were reported elsewhere [40]. Lowly expressed genes were filtered out, and the gene counts normalization was carried out by applying the variance stabilizing transformation (*VST*) function from DeSeq2 [41].

To identify significant gene-metabolite pairs, we analyzed the data considering two approaches, *i.e*., breed-specific (Duroc-Landrace) and FE-specific (low-high FE groups). Thus, we fitted a linear model for adjusting the read counts with the covariates using Limma R package [42]. For adjusting the data to identify breed-specific differences, we adopted the following model:

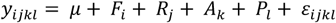

where,

*y*_*ijkl*_: is the normalized read counts;

μ: is the intercept;

*B*_*i*_: is the fixed effect of the FE group (two levels, high and low);

*R*_*j*_: is the covariate for the RIN values;

*A*_*k*_: is the covariate for the animal’s slaughter age, in days;

*P*_*l*_: is the fixed effect of the pen (8 levels);

*ε*_*ijkl*_: is the random residual effect associated with each observation.

To identify differences between high and low FE groups, the breed effect was added in the linear model, as follows:

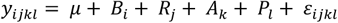

where,

*y*_*ijlk*_: is the normalized read counts;

μ: is the intercept;

*F*_*i*_: is the fixed effect of the breed (two levels, Duroc and Landrace);

*R*_*j*_: is the covariate for the RIN values;

*A*_*k*_: is the covariate for the animal’s slaughter age, in days;

*P*_*l*_: is the fixed effect of the pen (8 levels);

*ε*_*ijkl*_: is the random residual effect associated with each observation.

Regarding the identification of the metabolites, we used an untargeted metabolomic approach, as reported elsewhere [3]. In summary, 5 mL of blood samples at two-time points were collected from jugular venipuncture of each non-fasted pig into the EDTA tubes and immediately placed on ice. The plasma samples extracted from 109 animals (59 Duroc and 50 Landrace) were subjected to metabolomics analysis, as described in a previous study [3]. The metabolite data from this study were accessed using MetaboLights accession ID MTBLS1384 with a link: https://www.ebi.ac.uk/metabolights/MTBLS1384. Due to the need for paired data to carry the integrative analysis, only those samples with both metabolite and RNA-Seq data were used herein. The metabolite data from time-point 2 in 40 pigs were log-normalized before fitting into a linear model. Only those with the relative standard deviation > 0.15 were used based on the raw counts. As proposed for the RNA-seq, we adjusted the log-normalized metabolite data considering both approaches. First, the following model was employed for the breed-specific analysis:

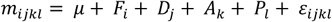

where,

*m*_*ijkl*_: is the is the log-normalized concentration of each metabolite (*n* = 749);

μ: is the intercept;

*F*_*i*_:: is the fixed effect of the FE group (two levels, high and low);

*D*_*j*_: is the fixed effect of the batch for metabolomic analysis (two levels);

*A*_*k*_: is the covariate for the sampling age, in days;

*P*_*l*_: is the fixed effect of the pen (8 levels);

*ε*_*ijkl*_: is the random residual effect associated with each observation.

For the FE-specific group approach, we fitted the data as follows:

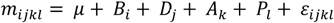

where,

*m*_*ijkl*_: is the is the log-normalized concentration of each metabolite (*n* = 749);

μ: is the intercept;

*F*_*i*_: is the fixed effect of the breed (two levels, Duroc and Landrace);

*D*_*j*_: is the fixed effect of the batch for metabolomic analysis (two levels);

*A*_*k*_: is the covariate for the sampling age, in days;

*P*_*l*_: is the fixed effect of the pen (8 levels);

*ε*_*ijkl*_: is the random residual effect associated with each observation.

The metabolites were annotated with HMDB (Human metabolome database) based on library search of the masses in HMDB with a mass uncertainty of 0.005Da or 5ppm. Those metabolites that did not correspond to HMDB entries were left unannotated and removed from the analysis.

### 4.3 Integration of transcriptomic and metabolomic data based on the linear model

To uncover the complex relationship between metabolites and genes, we adopted a linear model framework using the IntLIM (Integration of Linear model) R-package (version 0.1.0) (https://github.com/mathelab/IntLIM) [20]. The IntLIM approach allows us to integrate the metabolomic-transcriptomic data considering a case-control design. Thus, as initially proposed, we compared the breeds (Duroc *vs*. Landrace) and the FE groups (low and high FCR animals). As a quality control step from IntLIM, we filtered out genes with the lowest 5% of the variation. Gene and metabolite exploratory analyses were performed by applying Principal Component Analysis to identify breed- and FE-specific clusters.

The linear model for data integration is given as described in the following equation:

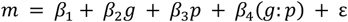

where,

*m*: is the log-normalized metabolite abundance;

*β*_1_: is the intercept;

*β*_2_*g*: is the normalized and adjusted gene expression level;

*β*_3_*P*: is the phenotype (FE group – high and low FE; or breed – Duroc and Landrace)

*β*_4_(*g*:*p*): is the interaction between gene expression and phenotype;

*ε*: is the residual effect associated with each observation (ε = N(0, σ))

A statistically significant two-tailed p-value of the gene-phenotype (g-p) interaction indicates the difference in the phenotype of FE groups (high and low) or breed (Duroc and Landrace) calculated by the slope relating gene-expression and metabolite abundance [20]. The two-tailed p-value indicates that the slope relating gene-expression and metabolite abundance is different from one phenotype compared to the other. Thus, it was used to identify gene-metabolite associations that are specific to a particular phenotype (breed – Duroc and Landrace, FE – low and high). We calculated the absolute difference in the Spearman correlation identified from IntLIM between the FE and breed groups to find the significant (p < 10^−7^) gene-metabolite pairs. The absolute difference between the FE group was estimated as (*r*_*High_cor*_ − *r*_*Low_cor*_) where (*r*_*High_cor*_) is the correlation given for a gene-metabolite pair in high-FE group (Table 2) while (*r*_*Low_cor*_) is the correlation given for a gene-metabolite pair in the low-FE group (Table 2). The absolute difference in the correlation between the breeds was estimated as (*r*_*Duroc_cor*_ − *r*_*Landrace_cor*_), where, (*r*_*Duroc_cor*_) is the correlation for a given gene-metabolite pair in Duroc (Table 1) while (*r*_*Landrace_cor*_) is the correlation for a gene-metabolite pair in Landrace (Table 1).

### 4.4 Pathway over-representation analysis

In breed-specific and FE-specific group, the same metabolite can be related to more than one gene and vice-versa. So, we screened for the common metabolites and genes in the group and referred them as unique metabolites and unique genes respectively in our study. We analyzed the unique metabolites in each group (breed-specific and FE-specific) using Metaboanalyst 4.0 (www.metaboanalyst.ca) [43]. We used three parameters for the pathway analysis: the pathway library, algorithm for pathway over-representation analysis, and algorithm for topological analysis. For our study, we selected the *Homo sapiens* (KEGG) pathway library to estimate the importance of the compound in a given metabolic pathway. For pathway over-representation and topology analysis, we used the hypergeometric test and relative-betweenness centrality algorithm, respectively, to measure the connections with the other nodes, including the number of shortest paths going through the node of interest.

Regarding the unique mapped (with chromosomal location information) genes, we carried out a co-functionality analysis using GeneMANIA [44] (www.genemania.org). GeneMANIA considers our query list of unique genes identified in each cluster (breed-specific and FE-specific) and allows us to predict the co-functional genes underlying similar functions. Thus, we analyzed the unique genes in each cluster (breed-specific – Duroc correlated cluster and Duroc anti-correlated cluster, FE-specific – High-FE correlated cluster) to identify the co-functional genes in each cluster. Next, we used the unique genes, as well as the co-functional genes in each cluster of each group, to identify the GO terms using GOrilla (Gene ontology enrichment analysis and visualization tool) (http://cbl-gorilla.cs.technion.ac.il/) [45]. To this end, the *Homo sapiens* were used as the reference, and the entire set of identified and annotated genes in our study (n = 15,187 genes) was used as a background.

## 5. Conclusions

This study applied a novel approach for metabolome-transcriptome integration study using the linear model unveiling potential gene-metabolite pairs affecting the biological processes related to FE in pigs. To the best of our knowledge, this is the first study to report the gene-metabolite pairs underlying FE in pigs. The approach followed here provided many interesting genes and metabolites with significant p-values. While some of the metabolites and genes identified were known with their association for FE, others are novel and provide new avenues for further research. The unique metabolites were associated with valine-leucine-isoleucine biosynthesis/degradation and arginine-proline metabolism. The unique genes enriched for sphingolipid catabolism and multicellular organismal process (breed-specific), and cGMP and purine metabolic processes (FE-specific) pathways. Further validation of genes, metabolites, and gene-metabolite interactions in larger cohorts will help us to elucidate the regulatory mechanisms affecting the pathways underlying FE.

## Supplementary Materials

Figure S1. The principal component analysis of metabolites and genes. A and B: PCA plot of genes and metabolites, respectively, in (1) Duroc and (2) Landrace; C and D: PCA plot of genes and metabolites, respectively, in high and low feed efficient groups.

Table S1. Pathways enriched by unique gene-metabolite pairs. Spreadsheet tabs are divided into a) metabolite enrichment analysis results by Metaboanalyst; b) list of the co-functional genes by GeneMania; c) gene ontology pathways enriched by Gorilla

## Author Contributions

HNK conceived and designed this “FeedOMICS” project, obtained funding as the main applicant. V.C and H.N.K designed blood sampling experiments, phenotype data collection, metabolite profiling, and RNA-Seq experiments. PB carried out biostatistical and bioinformatic data analysis and wrote the manuscript. All authors collaborated in the interpretation of results, discussion, and write up of the manuscript. All authors have read, reviewed, and approved the final manuscript.

## Acknowledgments

This study was funded by the Independent Research Fund Denmark (DFF) – Technology and Production (FTP) grant (grant number: 4184-00268B). VAOC received partial Ph.D. stipends from the University of Copenhagen and Technical University of Denmark. Authors thank Wellison Jarles Da Silva Diniz for helping in the interpretation of results. Authors thank SEGES – Pig Research Centre (VSP) Denmark for access to blood samples and phenotype datasets used in this study.

## Consent for publication

The blood sampling and experiment were approved and carried out in accordance with the Ministry of Environment and Food of Denmark, Animal Experiments Inspectorate under the license number (tilladelsesnummer) 2016-15-0201-01123, and C-permit granted to the principal investigator /senior author (HK).

## Availability of data and material

The metabolites dataset generated and analyzed during the current study is publicly available at the Metabolights database https://www.ebi.ac.uk/metabolights/MTBLS1384 with accession ID: MTBLS1384. (https://doi.org/10.1093/nar/gks1004; PubMed PMID: 2310955). The transcriptome dataset analyzed during the present study is also publicly available at the NCBI-GEO database with Accession ID GSE148889 and can be accessed at: https://www.ncbi.nlm.nih.gov/geo/query/acc.cgi?acc=GSE148889.

## References

1. Patience, J.F.; Rossoni-Serão, M.C.; Gutiérrez, N.A. A review of feed efficiency in swine: biology and application. J. Anim. Sci. Biotechnol. 2015, 6, 33, doi: 10.1186/s40104-015-0031-2.

2. He, B.; Li, T.; Wang, W.; Gao, H.; Bai, Y.; Zhang, S.; Zang, J.; Li, D.; Wang, J. Metabolic characteristics and nutrient utilization in high-feed-efficiency pigs selected using different feed conversion ratio models. Sci. China Life Sci. 2019, 62, 959–970, doi: 10.1007/s11427-018-9372-6.

3. Carmelo, V.A.O.; Banerjee, P.; da Silva Diniz, W.J.; Kadarmideen, H.N. Metabolomic networks and pathways associated with feed efficiency and related-traits in Duroc and Landrace pigs. Sci. Rep. 2020, 10, 1–14, doi: 10.1038/s41598-019-57182-4.

4. Godinho, R.M.; Bergsma, R.; Silva, F.F.; Sevillano, C.A.; Knol, E.F.; Lopes, M.S.; Lopes, P.S.; Bastiaansen, J.W.M.; Guimarães, S.E.F. Genetic correlations between feed efficiency traits, and growth performance and carcass traits in purebred and crossbred pigs. J. Anim. Sci. 2018, 96, 817–829, doi: 10.1093/jas/skx011.

5. Ren, P.; Yang, X.J.; Cui, S.Q.; Kim, J.S.; Menon, D.; Baidoo, S.K. Effects of different feeding levels during three short periods of gestation on gilt and litter performance, nutrient digestibility, and energy homeostasis in gilts. J. Anim. Sci. 2017, 95, 1232–1242, doi: 10.2527/jas2016.1208.

6. Do, D.N.; Strathe, A.B.; Ostersen, T.; Pant, S.D.; Kadarmideen, H.N. Genome-wide association and pathway analysis of feed efficiency in pigs reveal candidate genes and pathways for residual feed intake. Front. Genet. 2014, 5, 307, doi: 10.3389/fgene.2014.00307.

7. Novais, F.J.; Dromms, R.A.; Alexandre, P.A.; Pires, P.R.L.; Styczynski, M.P.-W.; Ferraz, J.B.S.; Fukumasu, H.; Iglesias, A.H. Identification of a metabolomic signature associated with feed efficiency in beef cattle. BMC Genomics 2019, 20, 1–10, doi: 10.1186/s12864-018-5406-2.

8. Rohart, F.; Paris, A.; Laurent, B.; Canlet, C.; Molina, J.; Mercat, M.J.; Tribout, T.; Muller, N.; Iannuccelli, N.; Villa-Vialaneix, N.; et al. Phenotypic prediction based on metabolomic data for growing pigs from three main european breeds. J. Anim. Sci. 2012, 90, 4729–4740, doi: 10.2527/jas.2012-5338.

9. D’Alessandro, A.; Marrocco, C.; Zolla, V.; D’Andrea, M.; Zolla, L. Meat quality of the longissimus lumborum muscle of Casertana and Large White pigs: Metabolomics and proteomics intertwined. J. Proteomics 2011, 75, 610–627, doi: 10.1016/j.jprot.2011.08.024.

10. Bertram, H.C.; Oksbjerg, N.; Young, J.F. NMR-based metabonomics reveals relationship between preslaughter exercise stress, the plasma metabolite profile at time of slaughter, and water-holding capacity in pigs. Meat Sci. 2010, 84, 108–113, doi: 10.1016/j.meatsci.2009.08.031.

11. Jing, L.; Hou, Y.; Wu, H.; Miao, Y.; Li, X.; Cao, J.; Michael Brameld, J.; Parr, T.; Zhao, S. Transcriptome analysis of mRNA and miRNA in skeletal muscle indicates an important network for differential Residual Feed Intake in pigs. Sci. Rep. 2015, 5, 11953, doi: 10.1038/srep11953.

12. Vincent, A.; Louveau, I.; Gondret, F.; Tréfeu, C.; Gilbert, H.; Lefaucheur, L. Divergent selection for residual feed intake affects the transcriptomic and proteomic profiles of pig skeletal muscle. J. Anim. Sci. 2015, 93, 2745–2758, doi: 10.2527/jas.2015-8928.

13. Horodyska, J.; Hamill, R.M.; Reyer, H.; Trakooljul, N.; Lawlor, P.G.; Mccormack, U.M.; Wimmers, K. RNA-Seq of Liver From Pigs Divergent in Feed Efficiency Highlights Shifts in Macronutrient Metabolism, Hepatic Growth and Immune Response. 2019, 10, doi: 10.3389/fgene.2019.00117.

14. Carrillo, J.A.; He, Y.; Li, Y.; Liu, J.; Erdman, R.A.; Sonstegard, T.S.; Song, J. Integrated metabolomic and transcriptome analyses reveal finishing forage affects metabolic pathways related to beef quality and animal welfare. Sci. Rep. 2016, 6, 25948, doi: 10.1038/srep25948.

15. Cavill, R.; Jennen, D.; Kleinjans, J.; Briedé, J.J. Transcriptomic and metabolomic data integration. Brief. Bioinform. 2016, 17, 891–901, doi: 10.1093/bib/bbv090.

16. Wilkinson, J.M. Re-defining efficiency of feed use by livestock. Animal 2011, 5, 1014–1022, doi: 10.1017/S175173111100005X.

17. Do, D.N.; Strathe, A.B.; Jensen, J.; Mark, T.; Kadarmideen, H.N. Genetic parameters for different measures of feed efficiency and related traits in boars of three pig breeds. J. Anim. Sci. 2013, 91, 4069–4079, doi: 10.2527/jas2012-6197.

18. Morales, P.E.; Bucarey, J.L.; Espinosa, A. Muscle Lipid Metabolism: Role of Lipid Droplets and Perilipins. J. Diabetes Res. 2017, 2017, 1–10, doi: 10.1155/2017/1789395.

19. Pedersen, B.K. Muscle as a Secretory Organ. In Comprehensive Physiology; John Wiley & Sons, Inc.: Hoboken, NJ, USA, 2013.

20. Siddiqui, J.K.; Baskin, E.; Liu, M.; Cantemir-Stone, C.Z.; Zhang, B.; Bonneville, R.; McElroy, J.P.; Coombes, K.R.; Mathé, E.A. IntLIM: integration using linear models of metabolomics and gene expression data. BMC Bioinformatics 2018, 19, 81, doi: 10.1186/s12859-018-2085-6.

21. Wang, M.; Zhang, X.; Kang, L.; Jiang, C.; Jiang, Y. Molecular characterization of porcine NECD, SNRPN and UBE3A genes and imprinting status in the skeletal muscle of neonate pigs. Mol. Biol. Rep. 2012, 39, 9415–9422, doi: 10.1007/s11033-012-1806-6.

22. Wahl, M.C.; Will, C.L.; Lührmann, R. The Spliceosome: Design Principles of a Dynamic RNP Machine. Cell 2009, 136, 701–718, doi: 10.1016/j.cell.2009.02.009.

23. Bottje, W.G.; Lassiter, K.; Piekarski-Welsher, A.; Dridi, S.; Reverter, A.; Hudson, N.J.; Kong, B.-W. Proteogenomics Reveals Enriched Ribosome Assembly and Protein Translation in Pectoralis major of High Feed Efficiency Pedigree Broiler Males. Front. Physiol. 2017, 8, doi: 10.3389/fphys.2017.00306.

24. Horodyska, J.; Reyer, H.; Wimmers, K.; Trakooljul, N.; Lawlor, P.G.; Hamill, R.M. Transcriptome analysis of adipose tissue from pigs divergent in feed efficiency reveals alteration in gene networks related to adipose growth, lipid metabolism, extracellular matrix, and immune response. Mol. Genet. Genomics 2019, 294, 395–408, doi: 10.1007/s00438-018-1515-5.

25. Mukiibi, R.; Vinsky, M.; Keogh, K.A.; Fitzsimmons, C.; Stothard, P.; Waters, S.M.; Li, C. Transcriptome analyses reveal reduced hepatic lipid synthesis and accumulation in more feed efficient beef cattle. Sci. Rep. 2018, 8, 7303, doi: 10.1038/s41598-018-25605-3.

26. Duan, Y.; Duan, Y.; Li, F.; Li, Y.; Guo, Q.; Ji, Y.; Tan, B.; Li, T.; Yin, Y. Effects of supplementation with branched-chain amino acids to low-protein diets on expression of genes related to lipid metabolism in skeletal muscle of growing pigs. Amino Acids 2016, 48, 2131–2144, doi: 10.1007/s00726-016-2223-2.

27. Reyer, H.; Oster, M.; Magowan, E.; Dannenberger, D.; Ponsuksili, S.; Wimmers, K. Strategies towards Improved Feed Efficiency in Pigs Comprise Molecular Shifts in Hepatic Lipid and Carbohydrate Metabolism. Int. J. Mol. Sci. 2017, 18, doi: 10.3390/ijms18081674.

28. Holeček, M. Effect of starvation on branched-chain α-keto acid dehydrogenase activity in rat heart and skeletal muscle. Physiol. Res. 2001, 50, 19–24.

29. Mukiibi, R.; Vinsky, M.; Keogh, K.; Fitzsimmons, C.; Stothard, P.; Waters, S.M.; Li, C. Liver transcriptome profiling of beef steers with divergent growth rate, feed intake, or metabolic body weight phenotypes1. J. Anim. Sci. 2019, 97, 4386–4404, doi: 10.1093/jas/skz315.

30. Sung, Y.J.; Pérusse, L.; Sarzynski, M.A.; Fornage, M.; Sidney, S.; Sternfeld, B.; Rice, T.; Terry, J.G.; Jacobs, D.R.; Katzmarzyk, P.; et al. Genome-wide association studies suggest sex-specific loci associated with abdominal and visceral fat. Int. J. Obes. 2016, 40, 662–674, doi: 10.1038/ijo.2015.217.

31. Zhang, L.; Rajbhandari, P.; Priest, C.; Sandhu, J.; Wu, X.; Temel, R.; Castrillo, A.; de Aguiar Vallim, T.Q.; Sallam, T.; Tontonoz, P. Inhibition of cholesterol biosynthesis through RNF145-dependent ubiquitination of SCAP. Elife 2017, 6, doi: 10.7554/eLife.28766.

32. Banerjee, P.; Carmelo, V.A.O.; Kadarmideen, H.N. Genome-Wide Epistatic Interaction Networks Affecting Feed Efficiency in Duroc and Landrace Pigs. Front. Genet. 2020, 11, doi: 10.3389/fgene.2020.00121.

33. Higgins, M.G.; Fitzsimons, C.; McClure, M.C.; McKenna, C.; Conroy, S.; Kenny, D.A.; McGee, M.; Waters, S.M.; Morris, D.W. GWAS and eQTL analysis identifies a SNP associated with both residual feed intake and GFRA2 expression in beef cattle. Sci. Rep. 2018, 8, 14301, doi: 10.1038/s41598-018-32374-6.

34. Pavlaki, N.; Nikolaev, V. Imaging of PDE2- and PDE3-Mediated cGMP-to-cAMP Cross-Talk in Cardiomyocytes. J. Cardiovasc. Dev. Dis. 2018, 5, 4, doi: 10.3390/jcdd5010004.

35. Valadkhan, S.; Gunawardane, L.S. Role of small nuclear RNAs in eukaryotic gene expression. Essays Biochem. 2013, 54, 79–90, doi: 10.1042/bse0540079.

36. Sun, J.S.; Manley, J.L. A novel U2-U6 snRNA structure is necessary for mammalian mRNA splicing. Genes Dev. 1995, 9, 843–854, doi: 10.1101/gad.9.7.843.

37. Ravi, S.; Schilder, R.J.; Kimball, S.R. Role of Precursor mRNA Splicing in Nutrient-Induced Alterations in Gene Expression and Metabolism. J. Nutr. 2015, 145, 841–846, doi: 10.3945/jn.114.203216.

38. Mansouri, E.; Khorsandi, L.; Moaiedi, M.Z. Grape seed proanthocyanidin extract improved some of biochemical parameters and antioxidant disturbances of red blood cells in diabetic rats. Iran. J. Pharm. Res. 2015, 14, 329–334, doi: 10.22037/ijpr.2015.1608.

39. Russell, J.R.; Sexten, W.J.; Kerley, M.S.; Hansen, S.L. Relationship between antioxidant capacity, oxidative stress, and feed efficiency in beef steers. J. Anim. Sci. 2016, 94, 2942–2953, doi: 10.2527/jas.2016-0271.

40. Victor AO. Carmelo, Kadarmideen, H.N. Genome regulation and gene interaction networks inferred from muscle transcriptome underlying feed efficiency in Pigs. 2020, 1–33, doi: https://doi.org/10.1101/2020.03.20.998203.

41. Anders, S.; Huber, W. Differential expression analysis for sequence count data. Genome Biol. 2010, 11, R106, doi: 10.1186/gb-2010-11-10-r106.

42. Ritchie, M.E.; Phipson, B.; Wu, D.; Hu, Y.; Law, C.W.; Shi, W.; Smyth, G.K. limma powers differential expression analyses for RNA-sequencing and microarray studies. Nucleic Acids Res. 2015, 43, e47–e47, doi: 10.1093/nar/gkv007.

43. Chong, J.; Soufan, O.; Li, C.; Caraus, I.; Li, S.; Bourque, G.; Wishart, D.S.; Xia, J. MetaboAnalyst 4.0: Towards more transparent and integrative metabolomics analysis. Nucleic Acids Res. 2018, 46, W486–W494, doi: 10.1093/nar/gky310.

44. Mostafavi, S.; Ray, D.; Warde-Farley, D.; Grouios, C.; Morris, Q. GeneMANIA: a real-time multiple association network integration algorithm for predicting gene function. Genome Biol. 2008, 9, S4, doi: 10.1186/gb-2008-9-s1-s4.

45. Eden, E.; Navon, R.; Steinfeld, I.; Lipson, D.; Yakhini, Z. GOrilla: a tool for discovery and visualization of enriched GO terms in ranked gene lists. BMC Bioinformatics 2009, 10, 48, doi: 10.1186/1471-2105-10-48.

