## Supplementary material for "Integrative Analysis of Metabolomic and Transcriptomic Profiles Uncovers Biological Mechanism of Feed Efficiency in Pigs": Figure S1

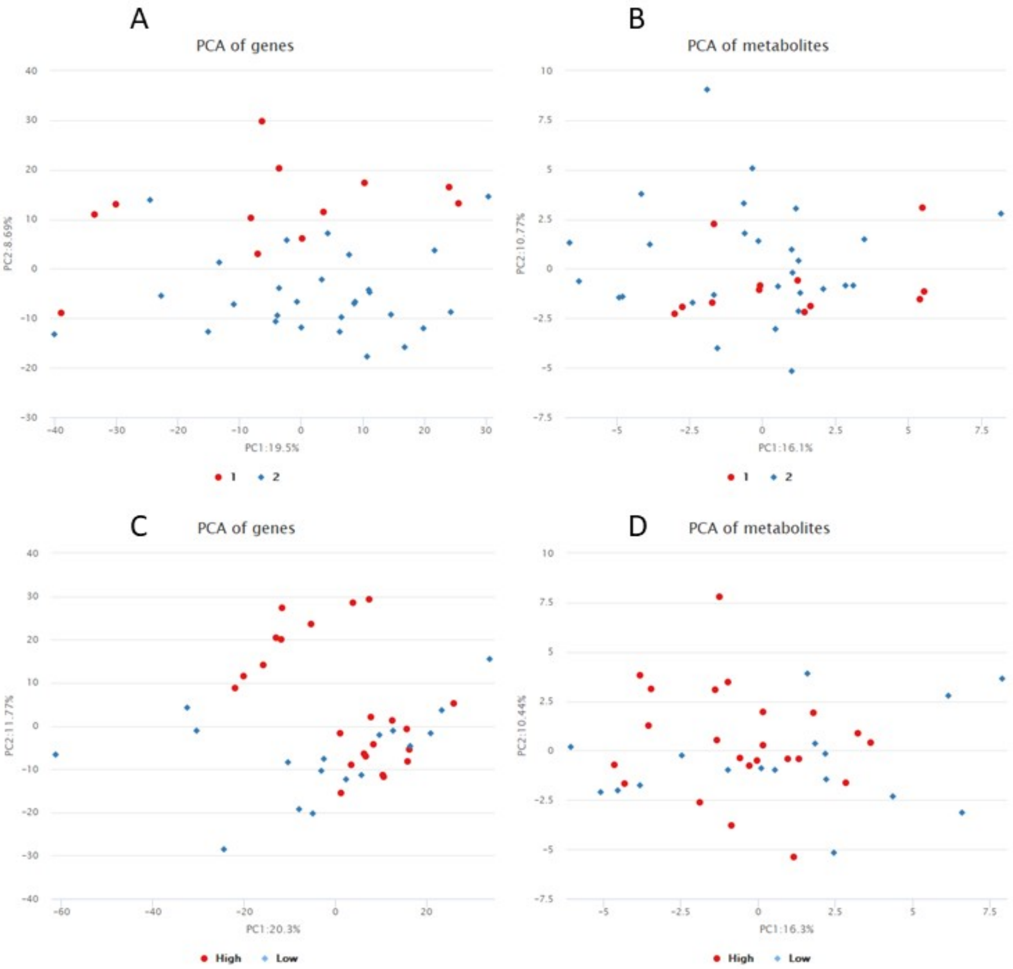


**Figure S1**. The principal component analysis of metabolites and genes. A and B: PCA plot of genes and metabolites, respectively, in (1) Duroc and (2) Landrace; C and D: PCA plot of genes and metabolites, respectively, in high and low feed efficient groups.
